# The Chromosome-Scale Genome of *Phyllanthus niruri* Reveals Candidate Genes and a Putative Biosynthetic Framework for Phyllanthin Formation

**DOI:** 10.64898/2026.05.15.725390

**Authors:** Kumari Khushi, Apoorva Ganesh, Aditya Sharma, Febina Ravindran, Subhashini Srinivasan, Bibha Chaudhary

## Abstract

*Phyllanthus niruri* (Phyllanthaceae) is a medicinally important herb known for producing phyllanthin, a bioactive dibenzylbutane lignan with reported hepatoprotective and antioxidant properties. However, the biosynthetic basis of phyllanthin production remains unresolved, largely due to the absence of a reference genome for the species.

We report a Chromosome-Scale Assembly of *P. niruri* generated by integrating PacBio HiFi long reads and Illumina short reads, followed by reference-guided scaffolding against *Phyllanthus cochinchinensis*. The assembly has an L50 of 7 and 97.6% BUSCO completeness.

Annotation predicted 19,254 protein-coding genes (91.1% functionally annotated), with phenylpropanoid biosynthesis emerging as the most enriched specialized-metabolism pathway in the genome.

Using pathway-guided genome mining, structural similarity analysis, and comparative metabolic reconstruction, we propose a putative biosynthetic pathway for phyllanthin originating from the phenylpropanoid-lignan branch through secoisolariciresinol-like intermediates, followed by terminal O-methylation reactions. A total of 305 unique candidate genes associated with the proposed pathway were identified, including expanded families of dirigent proteins, peroxidases, secoisolariciresinol dehydrogenases, and O-methyltransferases. Comparative transcriptomic analyses across related *Phyllanthus* species further supported the proposed pathway through coordinated expression of lignan-associated genes and tissue-specific enrichment of O-methyltransferases.

This work provides the first reference genome for *P. niruri* and a prioritized candidate gene set for functional characterization of phyllanthin biosynthesis.

## INTRODUCTION

*Phyllanthus niruri L*. (Phyllanthaceae), formerly classified under Euphorbiaceae, is a small annual herb (30–40 cm) distributed across the tropics, including the Amazon basin, South and Southeast Asia, and southern China. It is known by several vernacular names reflecting its ethnomedicinal prominence, including chanca piedra (“stone breaker”) in South America and bhumi amla in India (Kaur et al. 2017; Mediani et al. 2017).

*P. niruri* has been used in Ayurveda and other traditional systems for centuries, primarily as a hepatoprotective and lithotriptic agent, with documented use against jaundice, urolithiasis, hepatitis, and inflammatory disorders (Khatoon et al. 2006; Boim et al. 2010; Kaur et al. 2017).

More than 510 specialized metabolites have been reported across the genus *Phyllanthus*, with lignans, hydrolysable tannins, flavonoids, and triterpenoids representing the dominant chemical classes (Bagalkotkar et al. 2006; Patel et al. 2011; Kaur et al. 2017). Several *Phyllanthus* species are known to accumulate lignans such as phyllanthin, hypophyllanthin, and niranthin, along with tannins including corilagin, geraniin, and gallic acid (Ilangkovan et al. 2016; Kaur et al. 2017; Jantan et al. 2019). These metabolites have been associated with diverse pharmacological and biotechnological applications reported across the genus, including antiviral, hepatoprotective, antitumour, antidiabetic, antihypertensive, antimicrobial, antioxidant, and bioremediation-related activities (Bagalkotkar et al. 2006; Boim et al. 2010; Patel et al. 2011; Rizwan et al. 2021; Shanmugarajan et al. 2021; Verma et al. 2021). In addition, flavonoids such as quercetin- and kaempferol-derived glycosides have been widely characterized in *Phyllanthus* species and contribute substantially to their antioxidant potential (Falcone Ferreyra et al. 2012; Górniak et al. 2019; Liu et al. 2021).

Lignans are dimeric phenylpropanoids derived from the oxidative coupling of two monolignol units, classically coniferyl alcohol (Umezawa 2003). In the canonical lignan biosynthetic pathway, dirigent proteins mediate stereoselective coupling to produce (+)-pinoresinol (Davin et al. 1997), which is sequentially reduced by pinoresinol–lariciresinol reductase (PLR) to lariciresinol and subsequently secoisolariciresinol (Dinkova-Kostova et al. 1996). Secoisolariciresinol can either be oxidized by secoisolariciresinol dehydrogenase (SDH) to form matairesinol (Xia et al. 2001) or serve as a branch-point intermediate for the biosynthesis of diverse downstream dibenzylbutane lignans through tailoring reactions such as O-methylation, glycosylation, and ring modification. Phyllanthin is a dibenzylbutane lignan (Nawfetrias et al. 2024) structurally consistent with a tetra-O-methylated secoisolariciresinol-like scaffold; however, the genes and enzymes responsible for its biosynthesis in *Phyllanthus* have not yet been experimentally characterized.

Genomic resources for Phyllanthaceae have grown rapidly: chromosome-scale assemblies are now available for *P. cochinchinensis* (Zhang et al. 2022; Mahajan et al. 2023; Chen et al. 2024; Xin et al. 2026). These genomes have illuminated flavonoid, ascorbate, and lignin biosynthesis within the family. *P. niruri,* however, has lacked a reference genome, leaving its lignan-specific metabolism particularly phyllanthin biosynthesis, without a molecular framework.

In this study, we report a pseudochromosome-scale genome assembly of *P. niruri* generated by hybrid PacBio HiFi + Illumina sequencing and reference-guided scaffolding against *P. cochinchinensis* (Zhang et al. 2022). We characterize its repeat landscape, transcription factor repertoire, and specialized-metabolism gene content, and propose a putative biosynthetic route for phyllanthin grounded in genome-wide gene-family analysis and structural-similarity reasoning. In the absence of *P. niruri* transcriptome data, comparative expression analysis of candidate pathway genes was performed using publicly available transcriptomes from three related *Phyllanthus* species to evaluate whether the proposed biosynthetic pathway exhibits conserved transcriptional coherence across the genus.

## MATERIALS AND METHODS

### Contig-level assembly of *P. niruri*

Genome sequencing of *Phyllanthus niruri* was performed using both PacBio HiFi long-read and Illumina short-read platforms. To generate a high-quality draft assembly, PacBio HiFi reads were independently assembled using WTDBG2 (v2.5), Flye (v2.9.4-b1799), and Canu (v2.3-development). The resulting assemblies were merged using QuickMerge to obtain a consensus assembly. The merged assembly was further improved using Illumina short reads through MaSuRCA hybrid assembly. Redundant haplotigs and duplicated contigs were subsequently removed using the purge_dups pipeline to generate a non-redundant haploid representation of the genome.

### Chromosome-level assembly of *Phyllanthus cochinchinensis*

To support reference-guided scaffolding and comparative genomic analyses, the *P. cochinchinensis* genome was independently assembled from publicly available raw sequencing data (NCBI BioProject PRJNA825722). De novo assembly was performed using Flye, followed by Hi-C-based scaffolding using YAHS. Assembly statistics, including N50 and L50 values, were assessed using QUAST (v5.2.0), while genome completeness was evaluated using BUSCO v5.2.2 with the embryophyta_odb10 lineage dataset.

### Scaffold-level assembly of *P. niruri*

To evaluate genomic similarity between the two species, Illumina reads from *P. niruri* were mapped against the assembled *P. cochinchinensis* genome, resulting in an overall mapping rate of 48.52%. Based on this substantial sequence similarity, the *P. cochinchinensis* assembly was used as a reference for homology-guided scaffolding. The contig-level assembly of *P. niruri* was scaffolded using LRScaf (v1.1.10), followed by chromosome-level anchoring and scaffold correction using RagTag (v2.1.0) using the *P. cochinchinensis* genome as a reference for scaffold ordering and orientation.

### Comparative Genomics

Comparative genomic analyses were conducted to assess inter-species genome organization between *Phyllanthus niruri* and *Phyllanthus cochinchinensis*. Whole-genome alignments were performed using NUCmer (from MUMmer v4.0.0rc1) to identify conserved and rearranged genomic regions across the two assemblies. Syntenic relationships and large-scale structural variations were visualized using a custom Python script, enabling the identification of collinear blocks and structural rearrangements between the two species. Self-synteny analysis of the *P. niruri* genome was additionally carried out using NUCmer to detect intra-genomic duplications and paralogous regions, and the resulting alignments were similarly visualized using a custom Python script.

### Repeat Identification and Genome Masking

De novo identification of repetitive elements in the *P. niruri* genome was performed using RepeatModeler (v2.0.1). A custom repeat library was generated from the assembled genome using the integrated RepeatScout and RECON algorithms, along with LTR structural discovery implemented within the RepeatModeler pipeline.

The custom repeat library was subsequently used for genome masking and repeat classification using RepeatMasker, and the resulting annotation outputs were used to estimate repeat composition and genomic distribution of repetitive elements across the assembly.

### Gene Prediction

Gene prediction was performed on the soft-masked *P. niruri* genome assembly using AUGUSTUS (v3.5.0), an ab initio gene prediction tool. To improve prediction accuracy, AUGUSTUS was trained using conserved single-copy orthologs identified through BUSCO analysis with the embryophyta_odb10 dataset. The resulting species-specific training parameters were subsequently used for genome-wide gene prediction.

### Identification of Simple Sequence Repeats (SSRs)

Simple Sequence Repeats (SSRs) were identified in the *P. niruri* genome using the MIcroSAtellite Identification Tool (MISA) (https://github.com/cfljam/SSR_marker_design/blob/master/misa.pl). SSR mining was performed using default parameters to detect mono-, di-, tri-, tetra-, penta-, and hexanucleotide repeat motifs across the genome assembly.

### Transcription Factor Families Identification

Identification of transcription factor (TF) families in *Phyllanthus niruri* was performed using the predicted protein dataset generated from genome annotation. Protein sequences were queried against the Plant Transcription Factor Database (PlantTFDB/PlnTFDB v3.0) using BLASTP with an E-value cutoff of 1e−6.

### Functional Annotation

Functional annotation of the predicted protein-coding genes was performed using eggNOG-mapper v2 (Cantalapiedra et al. 2021), which assigns functional information based on orthology relationships. Predicted protein sequences obtained from AUGUSTUS gene prediction were queried against the eggNOG 5.0 database using the DIAMOND search mode for rapid and sensitive homology-based annotation.

Functional categories including Gene Ontology (GO) terms (Ashburner et al. 2000), Kyoto Encyclopedia of Genes and Genomes (KEGG) (Kanehisa and Goto 2000)pathway assignments, and Clusters of Orthologous Groups (COG) classifications were assigned based on the best orthologous matches identified through eggNOG-mapper.

### Identification and Analysis of Secondary Metabolic Pathway Genes

Genes involved in secondary metabolite biosynthesis were identified through pathway-guided functional annotation and comparative genomic analysis. Biosynthetic pathways associated with flavonoids, phenylpropanoids, lignans, terpenoids, alkaloids, and ascorbic acid metabolism were obtained from the KEGG database. Corresponding enzyme annotations and EC identifiers were manually curated and mapped to the annotated genomes of *Phyllanthus niruri* and *P. cochinchinensis* to identify pathway-associated candidate genes.

Additional metabolic information was retrieved from MetaCyc, BRENDA, and Rhea databases (Bansal et al. 2022) to identify genes potentially involved in genus-specific secondary metabolite biosynthesis. Particular emphasis was placed on the proposed phyllanthin biosynthetic pathway. Since the pathway has not been experimentally characterized, a hypothetical pathway model was constructed based on structural similarity to known lignan and phenylpropanoid-derived metabolites and their associated enzymatic reactions (Gang et al. 1999; Umezawa 2003). Candidate genes, including reductases, dehydrogenases, cytochrome P450s, and O-methyltransferases (OMTs), were identified using functional annotation, orthology analysis, and comparative transcriptomic evidence from related *Phyllanthus* species.

### Transcriptome Data Processing and Comparative Transcriptomic Analysis

RNA-Seq data for *P. niruri* are not currently available; we therefore used transcriptomes from three congeners as comparative evidence. Public RNA-Seq libraries were retrieved from the NCBI SRA for *P. cochinchinensis* (leaf: SRR18748705; stem: SRR18748704), *P. emblica* (PRJDB18024), and *P. amarus* (leaf: SRR1291302). Reads were quality- and adapter-trimmed using Trimmomatic (v0.39) and Cutadapt (v5.0), and quality-checked with FastQC. For each species, reads were pseudo-aligned with Kallisto (v0.48) against species-specific de novo transcriptome assemblies generated with Trinity (v2.15.2) and functionally annotated with eggNOG-mapper. Orthologous pathway genes across species were identified by reciprocal best BLAST hits against the *P. niruri* gene set, and expression values were Z-score normalized for visualization across tissues and species.

## RESULTS

### Genome Sequencing and Assembly

The final pseudochromosome-level assembly of *P. niruri* had a total size of 332 Mb with a GC content of 35.16%, scaffold N50 of 18.32 Mb, and L50 of 7 (Table 1). Genome completeness assessment using BUSCO with the embryophyta_odb10 dataset showed 97.6% completeness, indicating a highly complete assembly suitable for downstream comparative and functional genomic analyses.

**Table 1:** Comparative genomic features of *Phyllanthus niruri* and *Phyllanthus cochinchinensis*.

| Feature | *Phyllanthus niruri* | *Phyllanthus cochinchinensis* |
| --- | --- | --- |
| **Chromosome number (2n)** | 26 | 26 |
| **Assembly size** | 332 Mb FASTA | 284 Mb FASTA |
| **Scaffold N50** | 18.32 Mb | 10.32 Mb |
| **L50** | 7 | 7 |
| **GC content** | 35.16% | 35.4% |
| **BUSCO completeness** | 97.6% | 98.1% |
| **Predicted protein-coding genes** | 19,254 | 20,836 |
| **Functional annotation** | 91.1% | 98.01% |
| **Repeat content** | 54.72% | 59.12% |
| **LTR/Copia** | 21.73% | 22.91% |
| **LTR/Gypsy** | 14.27% | 11.94% |
| **DNA transposons** | 1.85% | — |
| **LINE elements** | 0.70% | 1.54% |

The *P. niruri* assembly showed comparable continuity and completeness to the *P. cochinchinensis* genome (Zhang et al. 2022) (Table 1). As chromosome-scale anchoring was performed using a reference-guided approach, the eleven major assembled scaffolds are referred to here as pseudochromosomes (Figure 1). Two additional smaller scaffolds corresponded to syntenic regions associated with *P. cochinchinensis* chromosomes 12 and 13 and likely represent partially unresolved assembly regions rather than differences in chromosome number between the two species.

**Figure 1.**
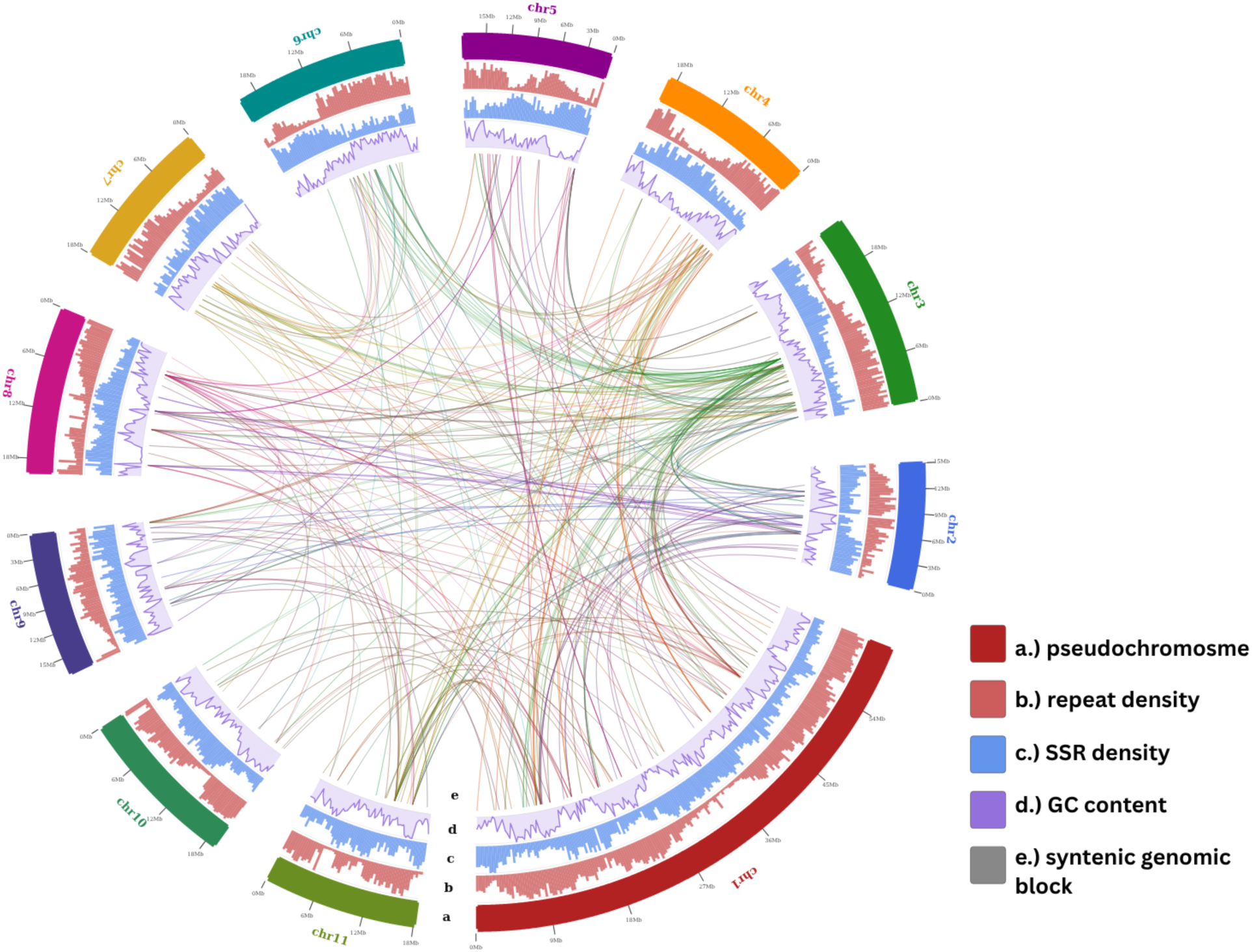
Circos representation of genomic features and syntenic relationships in the *Phyllanthus niruri* genome assembly. The outermost track represents the 11 pseudochromosomes of *P. niruri*. Inner tracks indicate genomic repeat density, simple sequence repeat (SSR) density, and GC content distribution across the genome. Colored links in the center represent syntenic genomic blocks identified within the assembly, illustrating large-scale duplicated and homologous regions among chromosomes. Genomic feature densities were calculated using fixed-size sliding windows across each pseudochromosome.

### Synteny Analysis

In order to perform whole-genome synteny analysis between *Phyllanthus niruri* and *Phyllanthus cochinchinensis* was performed using NUCmer implemented in MUMmer v4.0.0rc1, and syntenic relationships were visualized using custom Python scripts. Comparative genome alignment revealed strong macrosyntenic conservation across chromosomes 1–11 of both species, with extensive collinear blocks indicating a high degree of chromosomal conservation and genome structural similarity (Figure 2).

**Figure 2.**
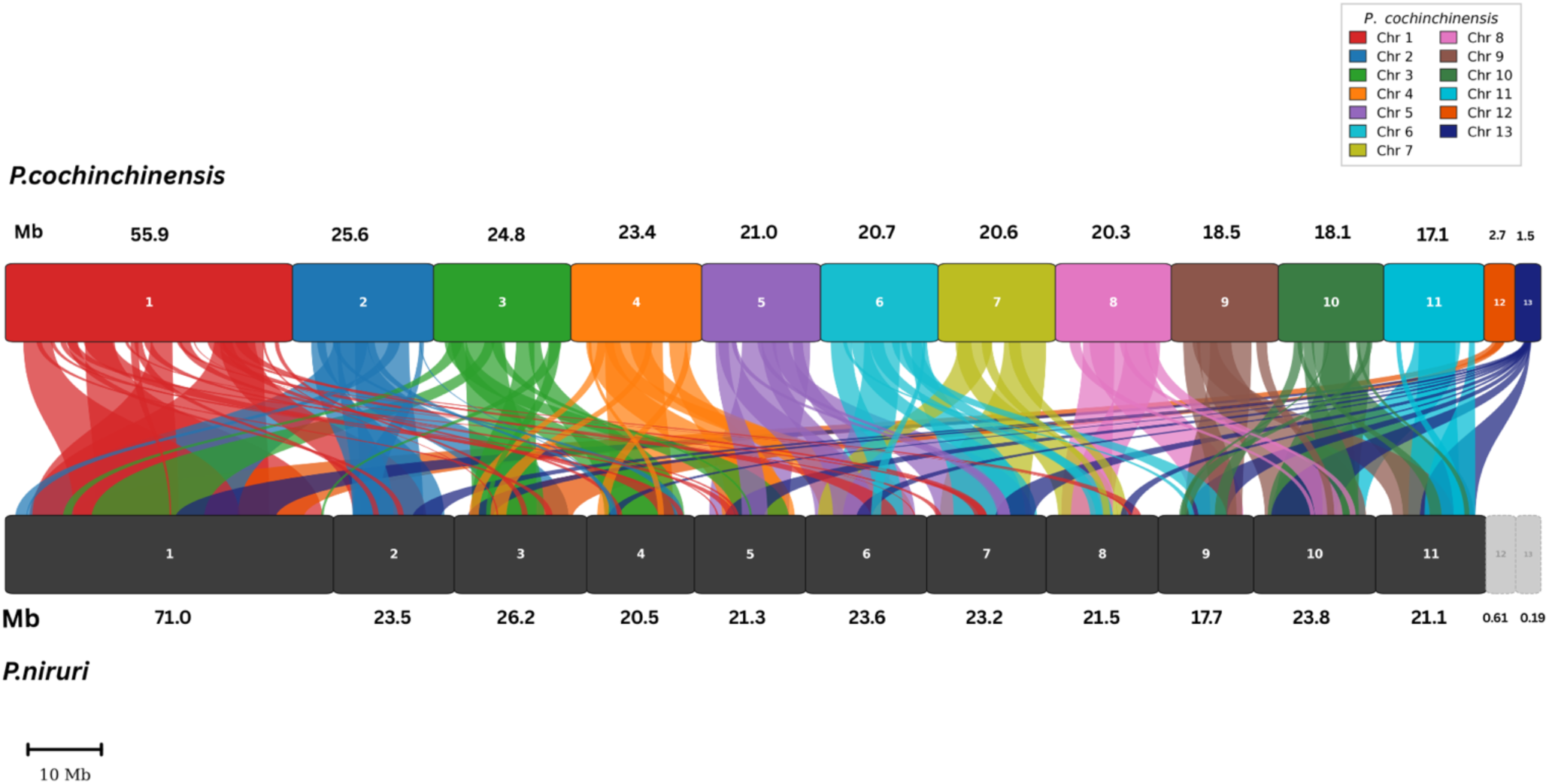
Whole-genome synteny analysis between *Phyllanthus niruri* and *Phyllanthus cochinchinensis*. Colored ribbons represent syntenic genomic blocks identified between corresponding chromosomes of the two species. Strong collinearity and extensive macrosyntenic conservation were observed across chromosomes 1–11, indicating highly conserved chromosomal organization within the genus *Phyllanthus*. The two smaller *P. niruri* scaffolds corresponding to chromosomes 12 and 13 showed fragmented syntenic associations distributed across multiple *P. cochinchinensis* chromosomes, suggesting partially unresolved assembly regions rather than true chromosomal reduction. Chromosome lengths are shown in megabases (Mb).

Karyotypic studies of *P. niruri* have reported a chromosome number of 2n = 26 with a basic chromosome number of x = 13 (Rahman et al. 2021), consistent with the chromosome organization observed in *P. cochinchinensis*. In the present assembly, *P. niruri* resolved into eleven major pseudochromosomes along with two comparatively smaller scaffolds (Figure 2). Targeted re-alignment of *P. cochinchinensis* chromosomes 12 and 13 against the complete *P. niruri* assembly identified syntenic regions distributed across multiple *P. niruri* pseudochromosomes rather than localized exclusively to the smaller scaffolds (Figure 2). These results indicate that orthologous genomic regions corresponding to chromosomes 12 and 13 are represented within the assembly, although some regions likely remain fragmented or distributed across partially anchored scaffolds. The conservation of syntenic blocks observed between the two species further supports a highly similar chromosomal organization within the genus *Phyllanthus*.

### Genome Annotation

Genome annotation for *Phyllanthus niruri* and *Phyllanthus cochinchinensis* was performed using AUGUSTUS with species-specific training generated from BUSCO-derived conserved orthologs. Gene prediction was carried out on soft-masked genome assemblies to identify protein-coding gene models and associated structural features.

A total of 19,254 protein-coding genes were predicted in the *P. niruri* genome, while 20,836 genes were identified in *P. cochinchinensis* (Zhang et al. 2022) (Table 1). Functional annotation of the predicted proteins was subsequently performed using EggNOG-mapper (Cantalapiedra et al. 2021) and BLASTP searches against the UniProt and *Arabidopsis thaliana* protein databases. In *P. niruri*, 17,538 genes (91.1%) were assigned putative functional annotations based on sequence homology and orthology relationships.

The annotated gene models provided a comprehensive framework for downstream comparative genomics, orthology inference, transcription factor identification, and secondary metabolite pathway analysis within the *Phyllanthus* genus.

### SSR Mining and Characterization

Genome-wide SSR mining identified 198,017 simple sequence repeats (SSRs) across the *Phyllanthus niruri* genome using the MISA pipeline. SSRs were distributed throughout all 11 pseudochromosomes, with chromosome 1 containing the highest number of SSRs (15,996), followed by chromosomes 8 and 3, while the remaining chromosomes showed relatively comparable SSR abundance (Figure 3a). SSR density varied across chromosomes, ranging from approximately 250–310 SSRs per Mb.

**Figure 3.**
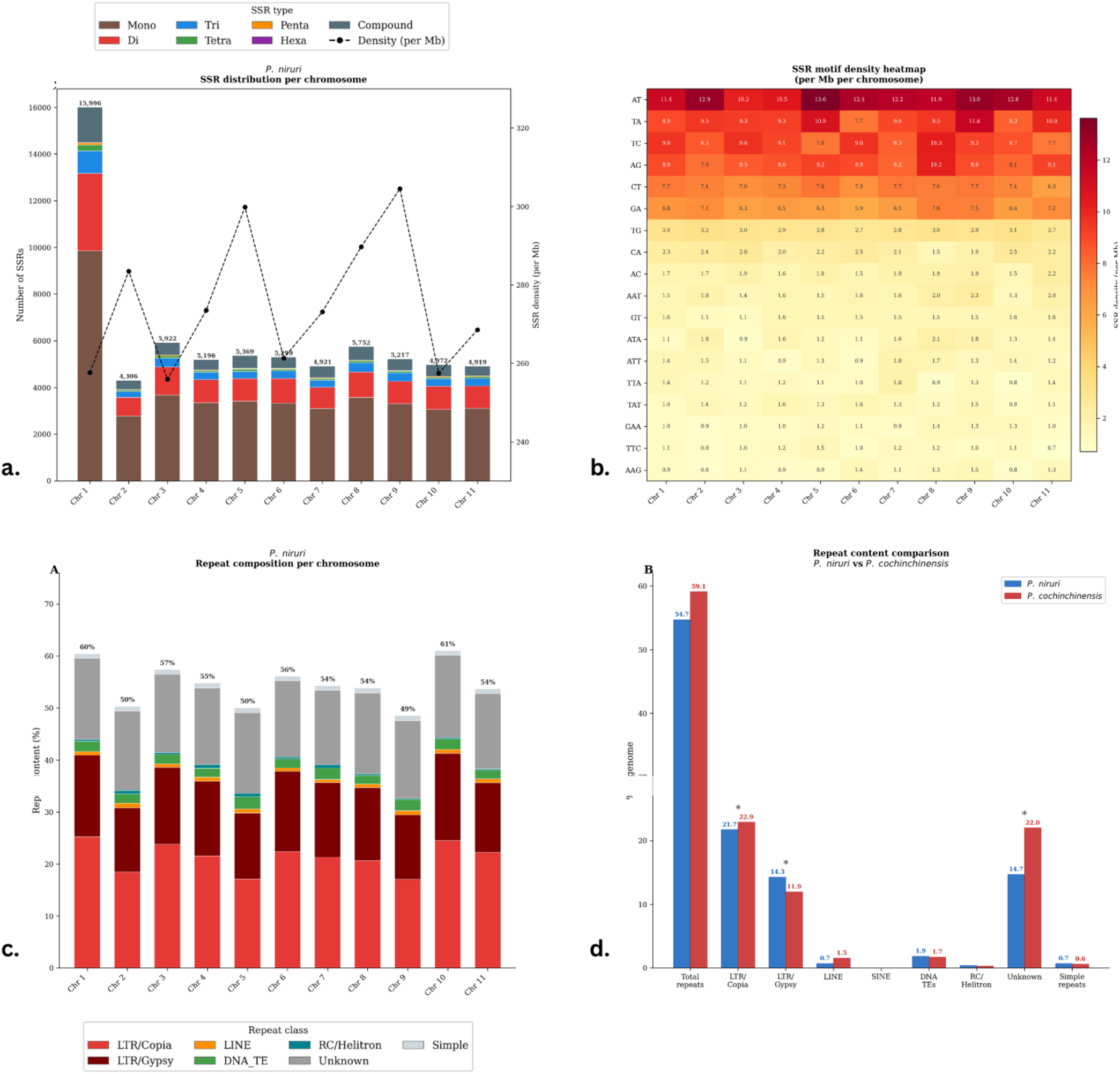
Genome-wide distribution of SSRs and repetitive elements in *Phyllanthus niruri*. (a) Chromosome-wise distribution and density of SSR classes identified across the 11 pseudochromosomes of *P. niruri*. Bars represent the abundance of mono-, di-, tri-, tetra-, penta-, hexa-, and compound SSRs, while the dashed line indicates SSR density per megabase. (b) Heatmap showing the density distribution of major SSR motifs across chromosomes. Warmer colors indicate higher motif abundance per megabase. AT- and TA-rich motifs were the most abundant SSR classes throughout the genome. (c) Chromosome-level repeat composition of the *P. niruri* genome. Stacked bars represent the proportion of major repeat classes including LTR/Copia, LTR/Gypsy, LINEs, DNA transposons, RC/Helitron elements, simple repeats, and unclassified repeats across pseudochromosomes. (d) Comparative repeat content analysis between *P. niruri* and *P. cochinchinensis*. The abundance of major repeat classes is shown as percentage genome coverage for each species, highlighting differences in LTR retrotransposon composition and unclassified repeat fractions.

Mononucleotide repeats represented the predominant SSR class, followed by dinucleotide and trinucleotide repeats, whereas tetra-, penta-, and hexanucleotide motifs occurred at substantially lower frequencies. Motif composition analysis revealed a strong predominance of AT-rich repeats throughout the genome. Among dinucleotide motifs, AT and TA repeats were the most abundant, while TC, AG, and CT motifs were also frequently represented (Figure 3b). Trinucleotide motifs including AAG, TTC, and GAA showed comparatively higher abundance than other trinucleotide repeat combinations.

The large number and diversity of SSR loci identified in this study provide a valuable genomic resource for future genetic diversity studies, molecular marker development, comparative genomics, and marker-assisted breeding applications within the Phyllanthus genus.

### Repeat Content Analysis

Repeat annotation using RepeatMasker revealed that repetitive elements occupy 54.72% of the *Phyllanthus niruri* genome, indicating a repeat-rich genome architecture comparable to other plant genomes. Comparative analysis showed slightly lower repeat content than the 59.12% reported for *P. cochinchinensis* (Zhang et al. 2022) (Figure 3d).

LTR retrotransposons represented the dominant repeat class in the *P. niruri* genome. Among these, LTR/Copia elements accounted for 21.73% of the genome and constituted the most abundant repeat category, followed by LTR/Gypsy elements at 14.27%. DNA transposable elements contributed 1.85% of the genome, whereas LINE and RC/Helitron elements represented comparatively smaller fractions. Unclassified repeats comprised 14.70% of the genome.

Chromosome-level repeat profiling demonstrated non-uniform repeat distribution across the 11 pseudochromosomes (Figure 3c). Chromosomes 10 and 1 exhibited the highest repeat densities, exceeding 60% repeat coverage, whereas the remaining chromosomes showed repeat content ranging between approximately 50–58%.

Comparative repeat landscape analysis between *P. niruri* and *P. cochinchinensis* revealed differences in repeat composition, particularly within LTR retrotransposon classes and unclassified repeats, reflecting ongoing repeat diversification within the Phyllanthus lineage (Zhang et al. 2022).

### Identification of Transcription Factor Families

Transcription factor (TF) analysis identified 3,148 putative TF genes in the *Phyllanthus niruri* genome, distributed across 57 TF families (Figure 4). Among the identified families, bHLH represented the most abundant class with 300 genes, followed by MYB-related (276), NAC (193), C2H2 zinc finger (186), ERF (170), MYB (164), and B3 (150) transcription factors. Other prominently represented families included FAR1, C3H, bZIP, WRKY, and M-type MADS. Several of the dominant TF families identified in *P. niruri* are associated with regulation of secondary metabolism, developmental processes, and stress-responsive signaling pathways. In particular, MYB and bHLH transcription factors are widely linked with phenylpropanoid and flavonoid biosynthesis (Falcone Ferreyra et al. 2012; Liu et al. 2021), while WRKY, NAC, AP2/ERF, and bZIP families are commonly involved in stress adaptation and specialized metabolite regulation.

**Figure 4.**
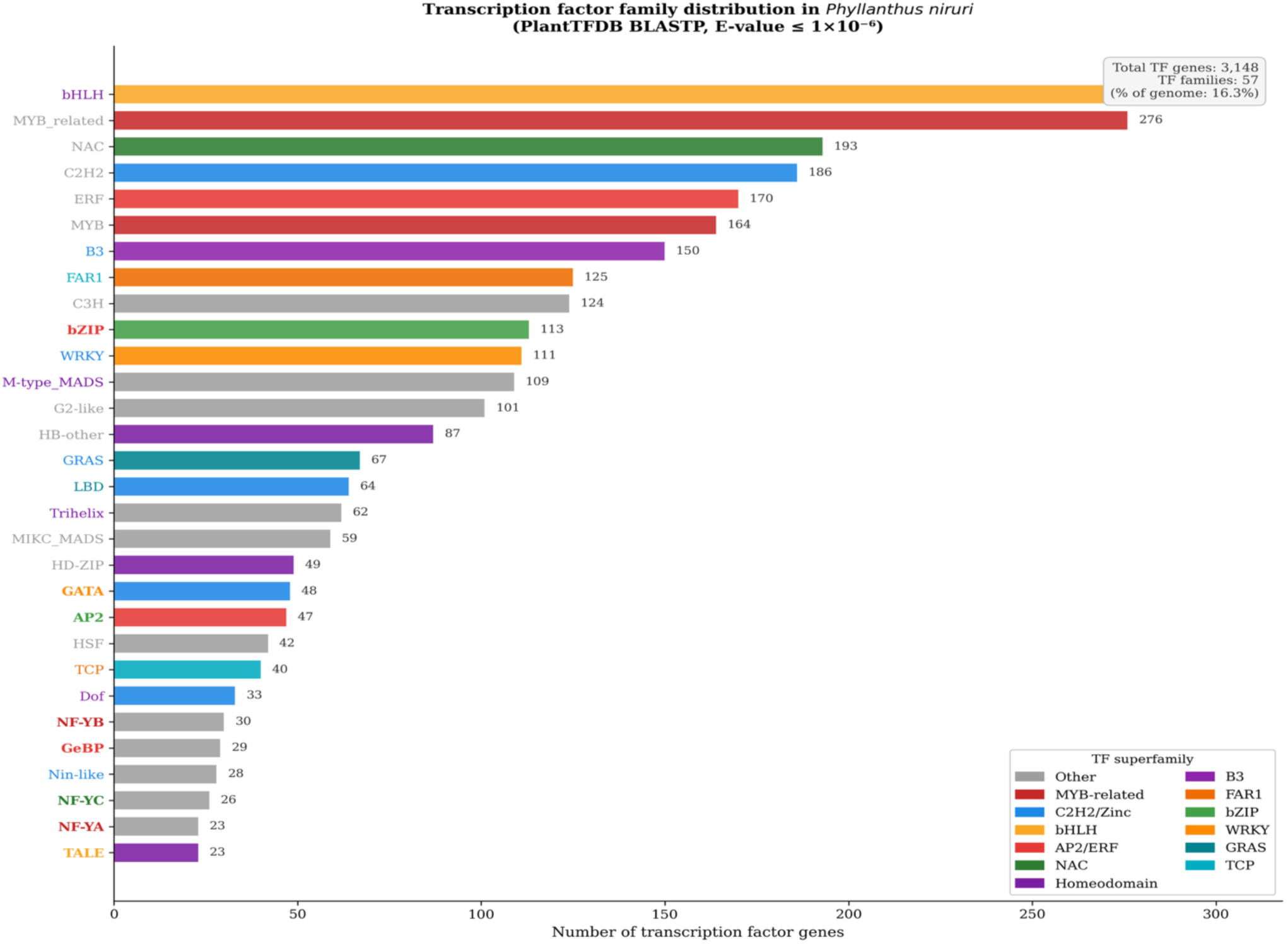
Distribution of transcription factor families identified in the *Phyllanthus niruri* genome. Predicted protein sequences were classified into transcription factor (TF) families based on BLASTP searches against the Plant Transcription Factor Database (PlantTFDB/PlnTFDB) using an E-value cutoff of 1 × 10⁻⁶. The plot shows the abundance of major TF families identified in the genome. A total of 3,148 TF genes belonging to 57 TF families were detected, representing approximately 16.3% of the predicted protein-coding genes. The bHLH, MYB-related, NAC, C2H2, ERF, and MYB families were among the most abundant transcription factor classes identified.

### Functional Annotation and Gene Ontology Classification

Functional annotation of predicted protein-coding genes in *Phyllanthus niruri* was performed using EggNOG-mapper based on orthology assignment and sequence similarity searches. Out of 19,254 predicted genes, 17,538 genes (91.1%) were successfully assigned putative functional annotations. Among these, 16,402 genes were further classified into COG functional categories, while a substantial proportion of genes were associated with Gene Ontology (GO) and KEGG pathway annotations (Kanehisa and Goto 2000).

GO classification recovered the expected distribution of functions for a plant genome of this size (Figure 5a). Within Biological Process, transcription regulation, transport, and metabolic processes were the largest categories; Molecular Function was dominated by catalytic activity and transcription factor activity; and Cellular Component was enriched for membrane-, plastid-, and nucleus-associated proteins.

**Figure 5.**
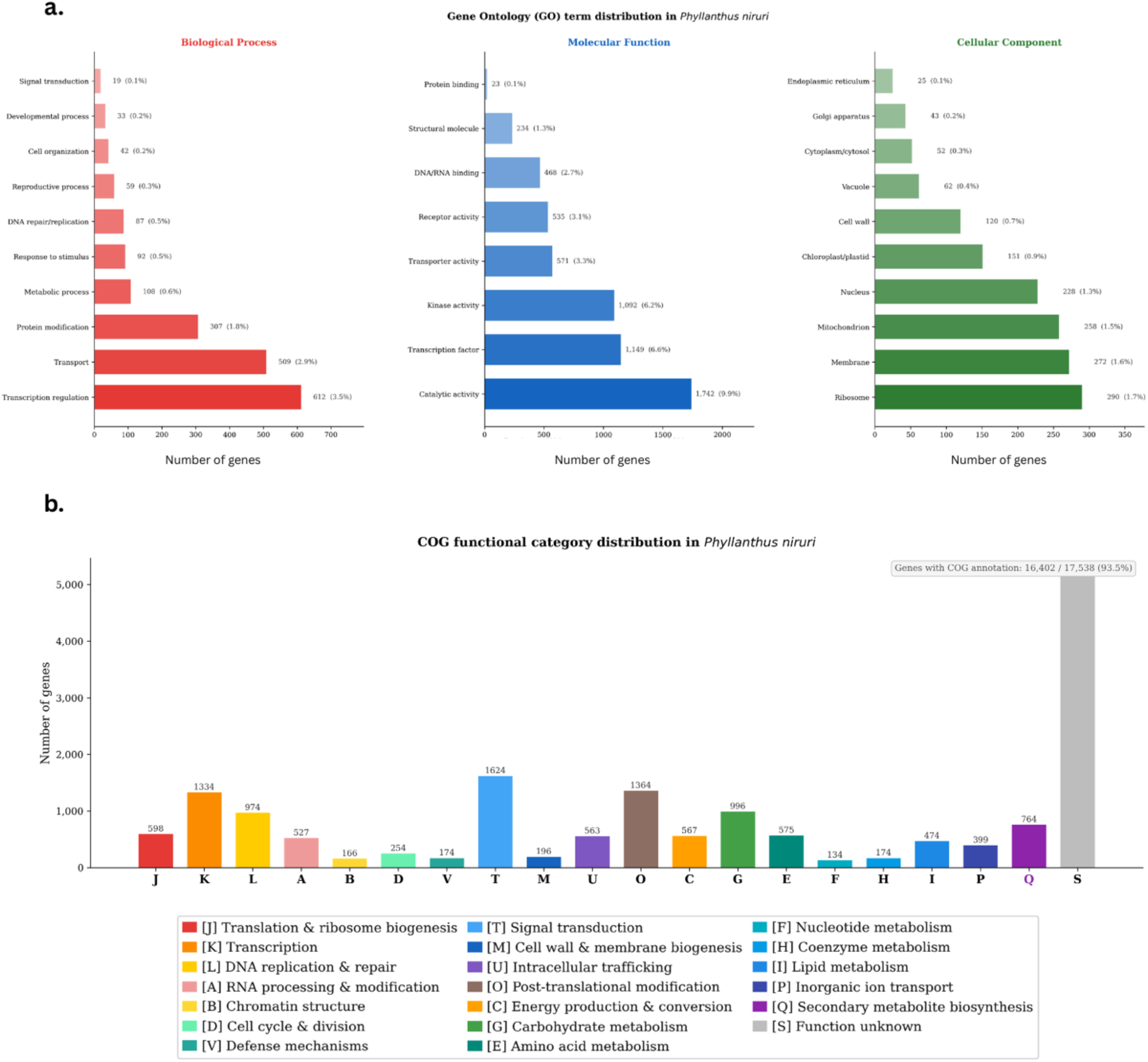
Functional annotation and Gene Ontology classification of predicted genes in *Phyllanthus niruri*. (a) Gene Ontology (GO) classification of annotated genes into the three principal ontologies: Biological Process, Molecular Function, and Cellular Component. Bars represent the number of genes assigned to each GO category. (b) Clusters of Orthologous Groups (COG) functional classification of annotated genes in *P. niruri*. Genes were categorized into major biological functions including signal transduction, transcription, metabolism, intracellular trafficking, defense mechanisms, and secondary metabolite biosynthesis.

COG functional classification revealed broad functional diversity within the *P. niruri* genome (Figure 5b). Genes involved in signal transduction mechanisms, post-translational modification, transcription, replication and repair, carbohydrate metabolism, intracellular trafficking, and secondary metabolite biosynthesis were highly represented. A substantial proportion of genes also belonged to the “function unknown” category, indicating the presence of potentially novel or poorly characterized genes within the genome.

### KEGG Pathway Annotation

KEGG pathway mapping assigned predicted genes in *Phyllanthus niruri* to a wide range of metabolic, signaling, and cellular pathways (Figure 6a). The most represented pathways included ribosome biogenesis, plant hormone signal transduction, plant–pathogen interaction, spliceosome, protein processing in the endoplasmic reticulum, and phenylpropanoid biosynthesis. Genes associated with purine metabolism, starch and sucrose metabolism, pyrimidine metabolism, and amino acid metabolism were also prominently represented, indicating extensive metabolic and regulatory complexity within the genome.

**Figure 6.**
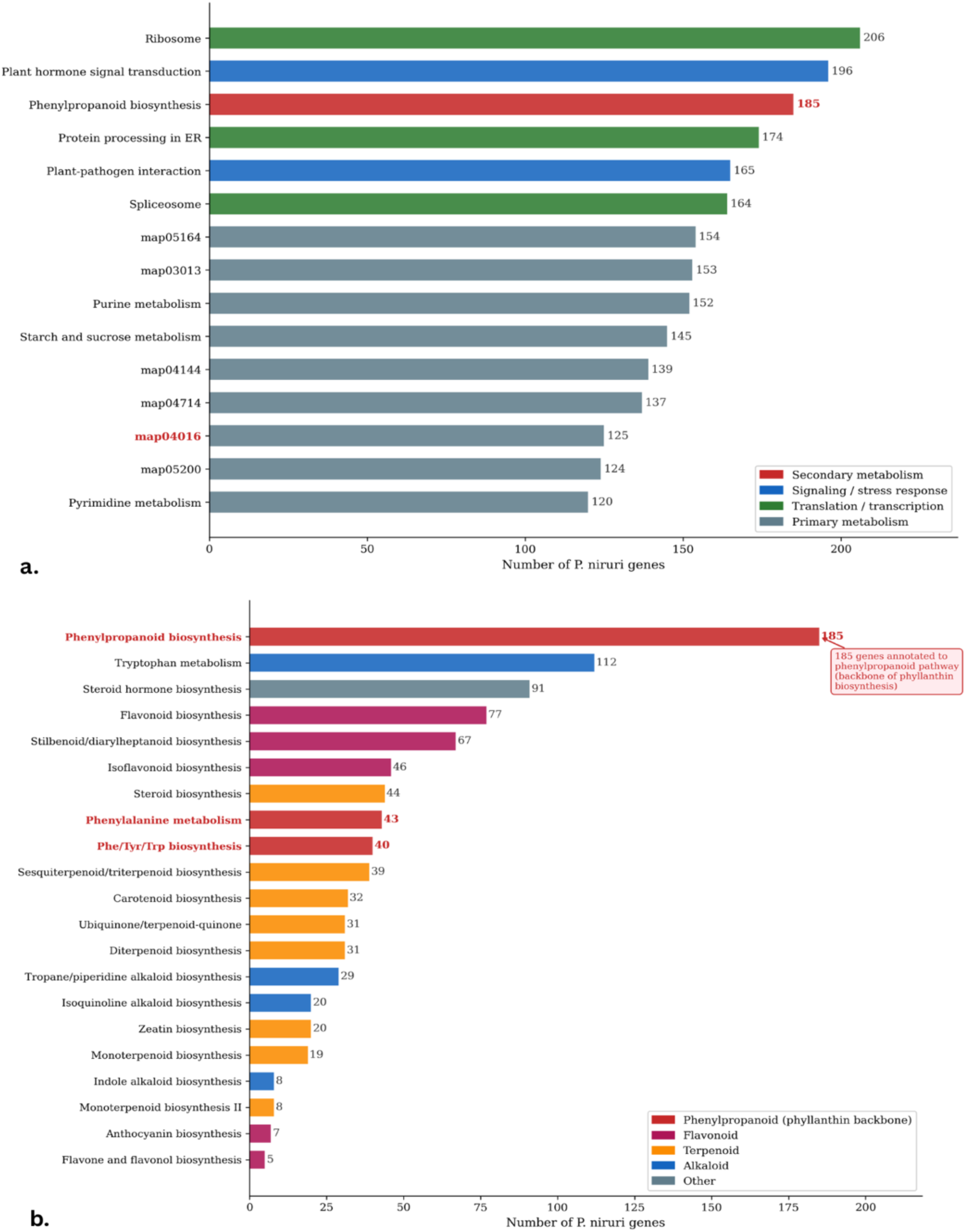
KEGG pathway enrichment and secondary metabolic pathway analysis in Phyllanthus niruri. (a) Major KEGG pathways identified in the *P. niruri* genome based on EggNOG and KEGG annotation. Pathways related to ribosome biogenesis, plant hormone signaling, phenylpropanoid biosynthesis, spliceosome function, and plant–pathogen interaction were among the most enriched categories. (b) Distribution of genes associated with major secondary metabolic pathways in *P. niruri*. Phenylpropanoid biosynthesis showed the highest enrichment, followed by flavonoid, stilbenoid, terpenoid, and alkaloid biosynthetic pathways, indicating extensive secondary metabolic capacity within the genome.

### Secondary Metabolic Pathway Analysis

Pathway-specific analysis identified extensive representation of genes associated with secondary metabolism (Figure 6b). Among these, phenylpropanoid biosynthesis represented the most enriched pathway, with 185 annotated genes identified in the *P. niruri* genome. Additional pathways associated with flavonoid, stilbenoid/diarylheptanoid, isoflavonoid, alkaloid, and terpenoid biosynthesis were also highly represented (Falcone Ferreyra et al. 2012; Liu et al. 2021).

Multiple terpenoid-associated pathways were detected, including sesquiterpenoid, diterpenoid, monoterpenoid, carotenoid, and ubiquinone biosynthesis. Genes associated with diverse alkaloid biosynthesis pathways, including tropane, piperidine, pyridine, isoquinoline, and indole alkaloid metabolism, were also identified. Collectively, these pathway annotations indicate broad biosynthetic capacity and extensive secondary metabolic diversity within the *P. niruri* genome

### Proposed Biosynthetic Pathway of Phyllanthin in *Phyllanthus niruri*

Functional annotation and pathway enrichment analysis identified extensive representation of genes associated with phenylpropanoid and lignan metabolism in the *Phyllanthus niruri* genome. KEGG pathway mapping revealed 185 genes associated with phenylpropanoid biosynthesis (Figure 6b), representing the most enriched secondary metabolite-associated pathway identified in the genome.

Although the complete biosynthetic pathway of phyllanthin remains unresolved, structural similarity analysis and comparative pathway reconstruction suggested that phyllanthin most likely originates from the phenylpropanoid-derived lignan branch through secoisolariciresinol-like intermediates (Umezawa 2003; Nawfetrias et al. 2024). Comparative analysis using KEGG, MetaCyc, and Rhea pathway databases identified lignan biosynthesis intermediates with strong structural resemblance to the phyllanthin backbone.

The proposed pathway proceeds from phenylalanine via the core phenylpropanoid route (PAL → C4H → 4CL → HCT/C3H → CCR → CAD) to coniferyl alcohol. Two coniferyl alcohol units undergo regio- and stereoselective oxidative coupling, mediated by class III peroxidases or laccases together with dirigent proteins, to form (+)-pinoresinol (Davin et al. 1997). (+)-Pinoresinol is sequentially reduced by pinoresinol–lariciresinol reductase (PLR) to (+)-lariciresinol and then to (−)-secoisolariciresinol (Dinkova-Kostova et al. 1996). In canonical lignan metabolism, secoisolariciresinol dehydrogenase (SDH) (Xia et al. 2001) further oxidizes secoisolariciresinol to matairesinol (Gang et al. 1999; Umezawa 2003); the abundance of SDH-family genes identified in *P. niruri* suggests this branch is active, but we propose that a parallel branch retains the dibenzylbutanediol scaffold and channels it into sequential O-methylation by SAM-dependent O-methyltransferases (OMTs), yielding the tetra-O-methylated phyllanthin structure (Figure 7).

**Figure 7.**
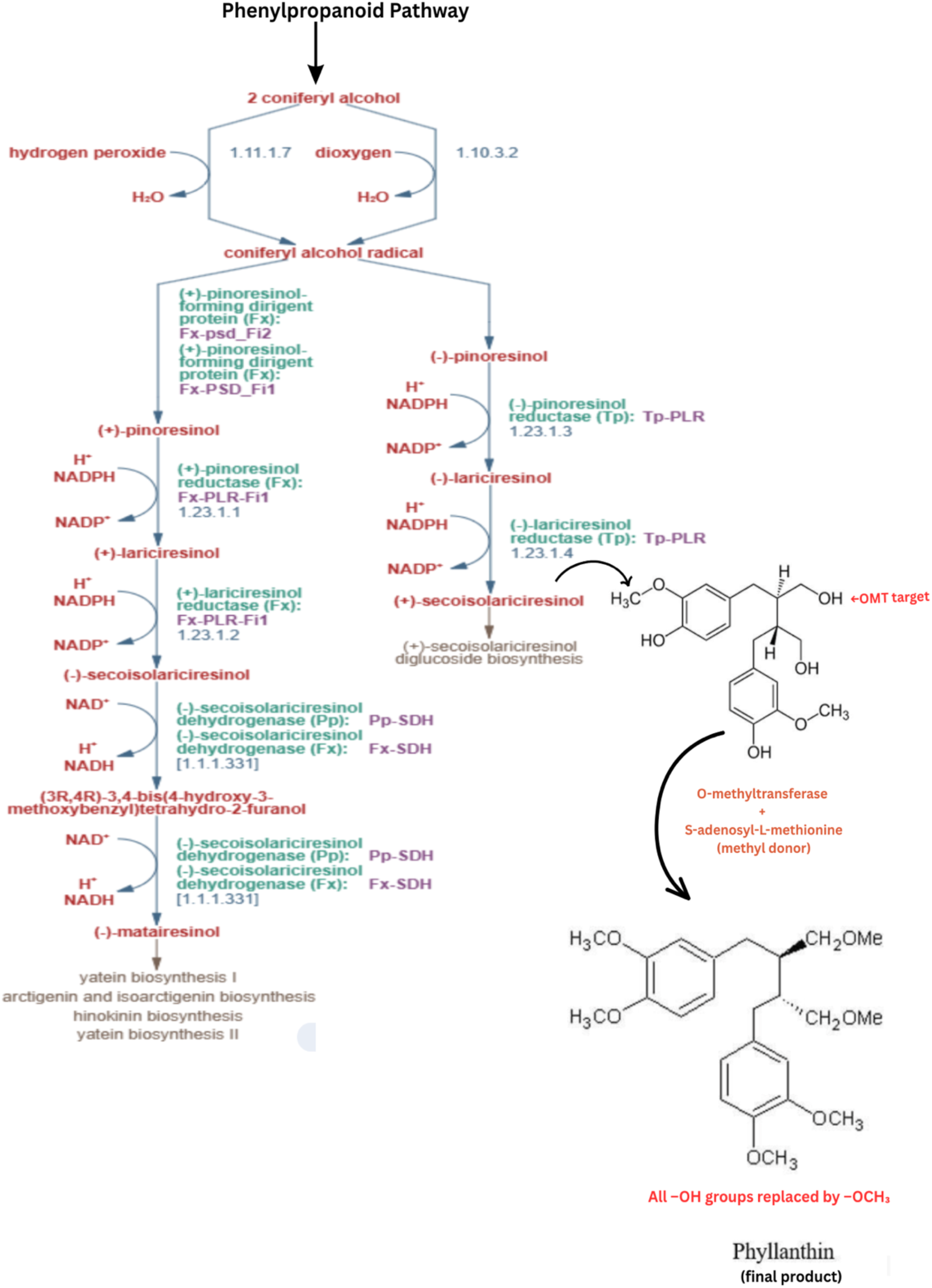
Proposed biosynthetic pathway of phyllanthin in *Phyllanthus niruri*. The pathway originates from the phenylpropanoid pathway through oxidative coupling of coniferyl alcohol molecules to form pinoresinol intermediates. Sequential reduction reactions mediated by pinoresinol/lariciresinol reductases (PLRs) and secoisolariciresinol dehydrogenases (SDHs) generate secoisolariciresinol-derived lignan intermediates. Based on structural similarity analysis, the hydroxylated lignan precursor is proposed to undergo multiple O-methylation reactions catalyzed by SAM-dependent O-methyltransferases (OMTs), resulting in the formation of phyllanthin. Arrows indicate the predicted enzymatic conversion steps inferred from KEGG lignan biosynthesis pathways and comparative genomic analyses.

Structural comparison between secoisolariciresinol-derived lignans and phyllanthin indicated that the primary distinction lies in replacement of hydroxyl (-OH) groups with methoxy (-OCH₃) substitutions. Based on this observation, the final biosynthetic steps were hypothesized to involve sequential O-methylation reactions catalyzed by SAM-dependent O-methyltransferases (OMTs), resulting in formation of the highly methoxylated phyllanthin structure (Figure 7).

Genome-wide pathway mining identified 376 pathway-associated gene assignments corresponding to 305 unique genes involved in the proposed phyllanthin biosynthesis pathway. These included PAL genes, dirigent proteins (Davin et al. 1997), SDH/LAR genes, peroxidases, and multiple expanded OMT families, supporting the presence of an extensive phenylpropanoid–lignan metabolic network in *P. niruri*.

### Comparative Expression Analysis of Candidate Phyllanthin Biosynthesis Genes

To further evaluate the proposed biosynthetic model, comparative expression analysis of candidate pathway genes was performed using publicly available transcriptome datasets from *Phyllanthus amarus*, *P. emblica*, and *P. cochinchinensis*.

Expression profiling of core phenylpropanoid pathway genes revealed elevated expression of TAL, PAL, C4H, 4CL, and COMT in *P. amarus* leaf tissues (Figure 8a), indicating active phenylpropanoid metabolism and lignan biosynthesis. In contrast, *P. emblica* exhibited comparatively lower expression levels across most pathway-associated genes.

**Figure 8.**
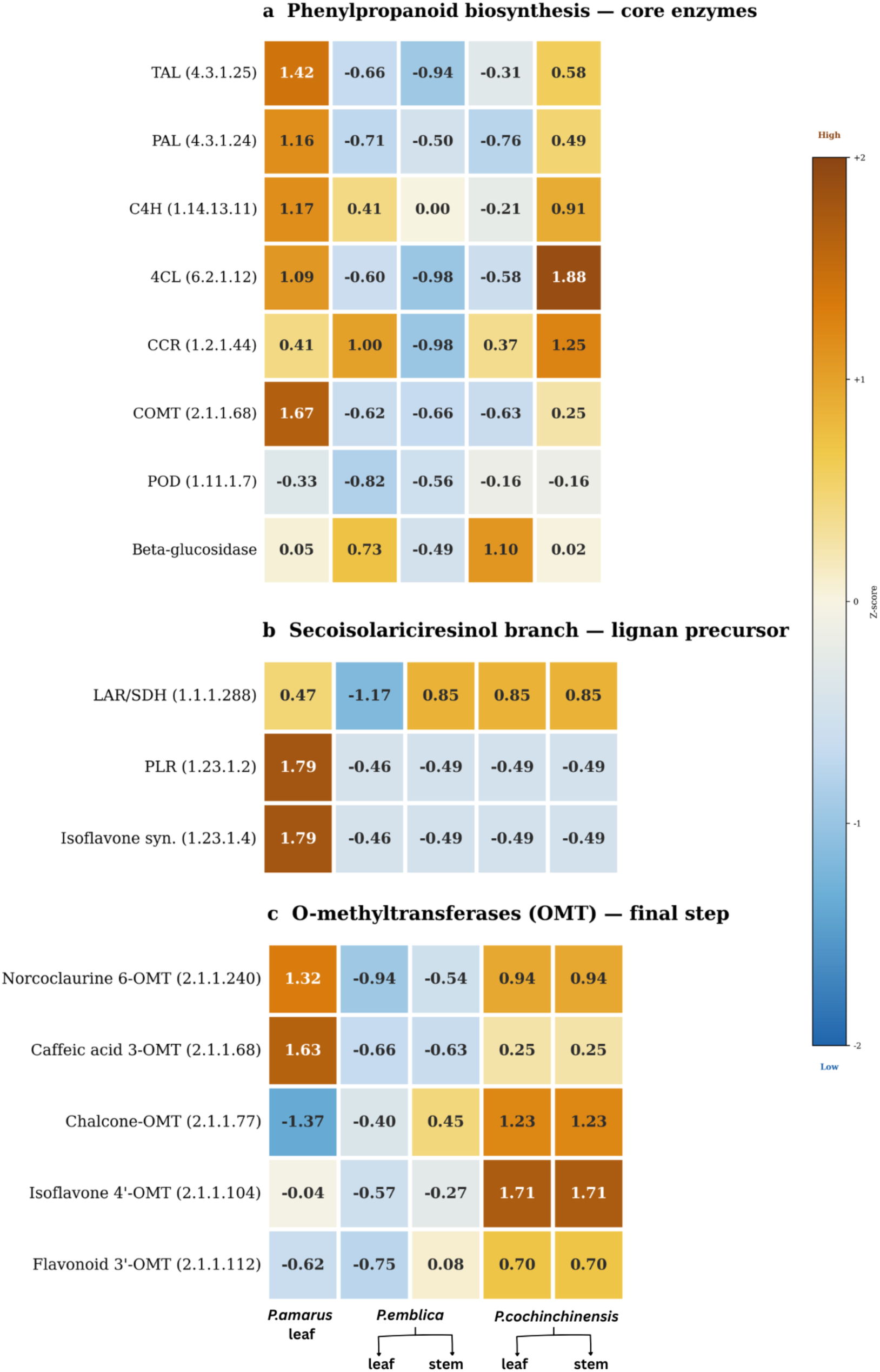
Comparative expression profiling of candidate phyllanthin biosynthesis genes across Phyllanthus species and tissues. Heatmap showing normalized expression (Z-score) of core phenylpropanoid pathway enzymes, including TAL, PAL, C4H, 4CL, CCR, COMT, POD, and β-glucosidase across transcriptomes of *P. amarus, P. emblica,* and *P. cochinchinensis.* (b) Expression profiles of enzymes associated with the secoisolariciresinol/lignan precursor branch, including pinoresinol/lariciresinol reductases (PLR) and secoisolariciresinol dehydrogenases (SDH/LAR). (c) Expression heatmap of candidate O-methyltransferases (OMTs) potentially involved in the terminal methylation reactions leading to phyllanthin biosynthesis, including norcoclaurine 6-O-methyltransferase, caffeic acid 3-O-methyltransferase, chalcone O-methyltransferase, isoflavone 4′-O-methyltransferase, and flavonoid 3′-O-methyltransferase. Expression values are represented as normalized Z-scores, where warmer colors indicate relatively higher expression and cooler colors indicate lower expression across tissues and species.

Genes associated with the proposed secoisolariciresinol branch, including PLRs and SDH/LAR enzymes, also showed increased expression in *P. amarus* leaf tissues (Figure 8b). Interestingly, several O-methyltransferase family genes, particularly norcoclaurine 6-O-methyltransferase, chalcone O-methyltransferase, and isoflavone 4′-O-methyltransferase, displayed strong enrichment in *P. cochinchinensis* stem tissues (Figure 8c).

Because O-methylation is proposed to represent the terminal step in phyllanthin biosynthesis (Figure 7), the elevated expression of OMT-associated genes in *P. cochinchinensis* stems suggests that downstream lignan modification and phyllanthin biosynthesis may preferentially occur in stem tissues in this species (Figure 8c). In contrast, pathway-associated genes in *P. amarus* were predominantly enriched in leaf tissues, indicating possible tissue-specific variation in lignan biosynthesis across *Phyllanthus* species.

Overall, the comparative expression profiles provide transcriptomic support for the proposed phyllanthin biosynthesis pathway and further support the hypothesis that phyllanthin formation proceeds through the phenylpropanoid–lignan branch followed by terminal O-methylation reactions.

## DISCUSSION

The present study provides the first high-quality chromosome-scale genome assembly of *Phyllanthus niruri*, an important medicinal species known for its diverse bioactive secondary metabolites and pharmacological properties. Despite its extensive ethnomedicinal use, limited genomic resources have hindered molecular investigations into therapeutically relevant compounds such as phyllanthin. Using a hybrid sequencing strategy we generated a chromosome-scale genome assembly enabling reliable characterization of repetitive regions, gene families, and secondary metabolite-associated loci. This genomic resource provides a foundation for comparative genomics, functional annotation, and investigation leading to specialized metabolism pathways within the genus *Phyllanthus*.

Comparative genomic analysis with *Phyllanthus cochinchinensis* revealed extensive macrosyntenic conservation across chromosomes 1–11 (Figure 2), indicating strong evolutionary conservation within the genus. In contrast, chromosomes 12 and 13 displayed comparatively fragmented syntenic relationships, suggesting possible lineage-specific rearrangements, chromosome fusion–fission events, or regions enriched in repetitive sequences (Figure 2). The overall conservation of chromosomal architecture supports a close evolutionary relationship between the two species while also highlighting genomic regions that may have contributed to species-specific metabolic diversification.

The chromosome-level genome assembly of *Phyllanthus cochinchinensis* (Zhang et al. 2022) indicated that the species shares the ancient core eudicot γ whole-genome duplication event with *Ricinus communis* and *Vitis vinifera*, with no evidence of a lineage-specific whole-genome duplication within *Phyllanthus*. Consistent with these findings, self-synteny analysis of the *P. niruri* genome did not reveal extensive duplicated chromosomal blocks indicative of recent polyploidization events (Figure 1). The expansion of gene families associated with lignan biosynthesis, including dirigent proteins, PALs, and O-methyltransferases, therefore likely resulted from tandem and dispersed duplication events rather than large-scale genome duplication.

Repetitive elements constituted a substantial proportion of the genome, with long terminal repeat retrotransposons representing the dominant class (Figure 3c). The dominance of LTR/Copia elements (21.73%) over LTR/Gypsy (14.27%) in *P. niruri* contrasts with the pattern in *P. cochinchinensis* (Figure 3d), where LTR/Copia and LTR/Gypsy are more nearly equal and may reflect lineage-specific transposon activity post-divergence (Zhang et al. 2022).

SSR analysis identified abundant microsatellite loci distributed throughout the *P. niruri* genome (Figure 3a,b), providing a useful resource for future applications in population genetics, phylogenetic analysis, conservation studies, and marker-assisted breeding within the genus *Phyllanthus*.

Functional annotation revealed strong enrichment of genes associated with catalytic activity, transcription regulation, signal transduction, intracellular trafficking, defense response, and membrane-associated functions, consistent with the metabolically active and specialized nature of medicinal plant genomes (Figure 5a,b) (Zhang et al. 2022; Mahajan et al. 2023)

KEGG pathway analysis (Kanehisa and Goto 2000) further revealed significant enrichment of phenylpropanoid biosynthesis (Figure 6b), along with flavonoid (Liu et al. 2021), stilbenoid, terpenoid, and alkaloid metabolism pathways. As phenylpropanoid metabolism serves as the precursor network for multiple bioactive compounds (Michal and Schomburg 2012), the enrichment of these pathways highlights the extensive phytochemical potential and medicinal relevance of *P. niruri*.

One of the major outcomes of this study was the reconstruction of a putative biosynthetic framework for phyllanthin formation. Although the upstream phenylpropanoid pathway is well characterized across plants, the downstream molecular steps leading specifically to phyllanthin biosynthesis have remained unresolved. Structural comparison of phyllanthin with known lignan intermediates revealed remarkable similarity with secoisolariciresinol-like compounds (Umezawa 2003; Nawfetrias et al. 2024), particularly in the conserved dibenzylbutane lignan backbone. Based on this observation, combined with pathway annotation and enzyme family identification, we hypothesized that phyllanthin biosynthesis proceeds through the phenylpropanoid–lignan branch via secoisolariciresinol-related intermediates followed by extensive O-methylation reactions (Figure 7).

Interestingly, metabolomic studies in *P. niruri* have consistently detected phyllanthin but rarely report detectable accumulation of secoisolariciresinol intermediates (Kaur et al. 2017; Mediani et al. 2017; Ruhela et al. 2023). This observation suggests that the intermediate may undergo rapid downstream conversion rather than accumulating as a stable metabolite. The high abundance of O-methyltransferase-associated genes identified in the genome strongly supports this possibility. In the proposed pathway, we hypothesize that hydroxyl groups present on secoisolariciresinol-like intermediates are sequentially methylated using S-adenosyl-L-methionine as a methyl donor, ultimately generating the highly methoxylated structure characteristic of phyllanthin.

Genome-wide screening identified 305 unique candidate genes associated with the proposed pathway, including enzymes involved in core phenylpropanoid metabolism, lignan formation, oxidative coupling, and terminal methylation reactions. The disproportionate expansion of dirigent proteins and OMTs relative to upstream phenylpropanoid genes may suggests that lignan diversification particularly stereoselective coupling and tailoring has been the dominant locus of selection in the Phyllanthus lignan pathway, rather than precursor supply (Figure 6b).

To further assess the proposed biosynthetic framework, comparative transcriptomic analyses were performed using publicly available datasets from *P. amarus* (Bose Mazumdar and Chattopadhyay 2016), *P. emblica* (Mahajan et al. 2023), and *P. cochinchinensis* (Zhang et al. 2022). As these species have previously been reported to accumulate phyllanthin or related lignans (Patel et al. 2011; Ilangkovan et al. 2016; Abd Rani et al. 2021), they provided suitable systems for comparative validation of pathway-associated genes.

Expression profiles showed coordinated expression of phenylpropanoid-, lignan-, and O-methyltransferase-associated genes across species and tissues, supporting conservation of the proposed biosynthetic framework (Figure 7). *P. amarus* leaf tissues exhibited relatively strong expression of core pathway genes, consistent with reports describing higher lignan accumulation in leaves (Patel et al. 2011; Ilangkovan et al. 2016), whereas *P. emblica* showed comparatively lower expression of several pathway-associated genes (Mahajan et al. 2023). Interestingly, *P. cochinchinensis* displayed elevated expression of downstream O-methyltransferases in stem tissues, suggesting possible tissue-specific partitioning of terminal lignan modification and phyllanthin biosynthesis within the genus (Sun et al. 2023).

This study provides a genomic framework for investigating phyllanthin biosynthesis in *Phyllanthus niruri*; however, experimental validation of the proposed pathway remains necessary. Future transcriptomic, metabolomic, and functional analyses of candidate lignan-associated genes will help refine the biosynthetic model and improve understanding of tissue-specific regulation of phyllanthin formation.

More broadly, the integration of chromosome-scale genomics with comparative and pathway-guided analyses demonstrates an effective strategy for resolving specialized metabolic pathways in medicinal plants. The candidate genes identified here may further support future synthetic biology and metabolic engineering approaches for scalable production of phyllanthin and related lignans.

## CONCLUSION

This study presents the first chromosome-scale reference genome of *Phyllanthus niruri*, providing a comprehensive genomic resource for investigating the molecular basis of specialized metabolism in this medicinally important species. The high-quality assembly enabled detailed characterization of genome organization, repeat architecture, transcription factor diversity, and secondary metabolite-associated pathways. Comparative genomic analyses revealed strong chromosomal conservation within the genus *Phyllanthus*, while functional annotation highlighted extensive enrichment of phenylpropanoid and lignan biosynthesis pathways. Through integrative genome mining, structural similarity analysis, and comparative transcriptomic evidence, we propose a putative biosynthetic framework for phyllanthin formation involving secoisolariciresinol-like intermediates followed by terminal O-methylation reactions. The identification of expanded lignan-associated gene families and coordinated expression of candidate pathway genes across related *Phyllanthus* species further supports the proposed pathway model. Together, these findings establish an important foundation for future functional validation of phyllanthin biosynthesis, comparative evolutionary studies, metabolic engineering, and molecular breeding of medicinal *Phyllanthus* species, while also helping bridge traditional ethnobotanical knowledge with modern plant genomics.

## DATA AVAILABILITY

Available on Request.

## ACKNOWLEDGEMENTS

PacBio HiFi reads were generated using services from Nucleome Informatics. The authors thank Government of Karnataka for funding for sequencing and data analysis personnel via a BioIT grant and computing infrastructure via the Department of Information Technology, Biotechnology and Science and Technology. The authors also acknowledge using computing resources obtained as part of DBT Builder Sanction no. BT/INF/22/SP45402/2022 dated March 8, 2022, CCB Sanction no. BT/PR40212/BTIS/137/40/2022 dated December 19, 2022, and DST-FIST Sanction no. SR/FST/LSI-536/2012.

